# Population analysis of *Vibrio cholerae* in aquatic reservoirs reveals a novel sister species *(Vibrio paracholerae* sp. nov.) with a history of association with human infections

**DOI:** 10.1101/2021.05.05.442690

**Authors:** Mohammad Tarequl Islam, Tania Nasreen, Paul Kirchberger, Kevin Y. H. Liang, Fabini Orata, Fatema-Tuz Johura, Monica S. Im, Cheryl L. Tarr, Munirul Alam, Yann F. Boucher

## Abstract

Most efforts to understand the biology of *Vibrio cholerae* have focused on a single group, the pandemic-generating lineage harbouring the strains responsible for all known cholera pandemics. Consequently, little is known about the diversity of this species in its native aquatic environment. To understand the differences in the *V. cholerae* populations inhabiting in regions with varying history of cholera cases and how that might influence the abundance of pandemic strains, a comparative analysis of population composition was performed. Little overlap was found in lineage compositions between those in Dhaka (cholera endemic) located in the Ganges delta, and of Falmouth (no known history of cholera), a small coastal town on the US East Coast. The most striking difference was the presence of a group of related lineages at high abundance in Dhaka which was completely absent from Falmouth. Phylogenomic analysis revealed that these lineages form a cluster at the base of the phylogeny of *V. cholerae* species, sufficiently differentiated genetically and phenotypically to form a novel species. Strains from this species have been anecdotally isolated from around the world and were isolated as early as 1916 from a British soldier in Egypt suffering from choleraic diarrhoea. In 1935 Gardner and Venkatraman unofficially referred to a member of this group as *Vibrio paracholerae*. In recognition of this earlier designation, we propose the name *Vibrio paracholerae*, sp. nov. for this bacterium. Genomic analysis suggests a link with human populations for this novel species and substantial interaction with its better-known sister species.

**Importance:** Cholera continues to remain a major public health threat around the globe. Understanding the ecology, evolution and environmental adaptation of the causative agent *Vibrio cholerae* and tracking the emergence of novel lineages with pathogenic potential are essential to combat the problem. In this study, we investigated the population dynamics of *Vibrio cholerae* in an inland locality which is known as endemic for cholera and compared with that of a cholera free coastal location. We found the consistent presence of the pandemic generating *V. cholerae* in cholera-endemic Dhaka and an exclusive presence of a lineage phylogenetically distinct from other *V. cholerae*. Our study suggests that this lineage represents a novel species having pathogenic potential and a human link to its environmental abundance. The possible association with human population, co-existence and interaction with toxigenic *V. cholerae* in the natural environment make this potential human pathogen an important subject for future studies.

## Introduction

*Vibrio cholerae* is the causative agent of cholera, the disease which has shaken human civilization from the last few centuries and continues to be a public health threat, especially to the developing world (1, 2). Its pathogenesis and epidemiology have been extensively studied, but the aquatic part of its life cycle is still not fully understood. Even though *V. cholerae* is a well defined, model species for microbial ecology research, strikingly, few close relatives have been found for this species in recent years, most being initially classified as *V. cholerae*-like bacteria. One of them was occasional human pathogen *Vibrio mimicus*, which was proposed as a new species in 1981 based on phenotypic characteristics (3). Later, genome-based studies established the molecular basis of its importance as a pathogen, close association and exchange of important virulence genes with *V. cholerae* (4, 5). Two other closely related novel species, *Vibrio parilis* and *Vibrio metoecus*, were more recently isolated alongside *V. cholerae* from coastal waters (6, 7) and found to exchange genetic material with *V. cholerae* in aquatic environments (6, 8). Biological information on the close relatives of a dangerous environmental pathogen like *V. cholerae* is of significance, because of their potential as emerging pathogens themselves and their interaction with *V. cholerae* in its natural habitats. Even though this diverse species is ubiquitous in tropical and temperate coastal waters world-wide, cholera is only caused by a specific lineage of *Vibrio cholerae*, in which the O1 antigen is ancestral (9, 10). It is not clear whether aquatic *V. cholerae* maintains a significantly different population structure in cholera endemic and non-endemic areas, and if this structure is influenced by co-occurring species. This is a crucial gap in our understanding of the factors defining cholera endemicity and driving local and global biogeographic dispersal patterns of *V. cholerae*. It has recently become possible to investigate the details of the population structure of *V. cholerae* and its close relatives, using a molecular marker based on a single copy housekeeping gene (*viuB*, vibriobactin utilization protein subunit B), which provides subspecies level resolution (11). This method was used to study a cholera-free region on the east coast of the USA, the Oyster Pond ecosystem (Falmouth, USA), where differences in abundance of individual alleles in particular locations/habitats indicated potential adaptation to ecological conditions at the subspecies level (11, 12). A similar study was performed in *V. cholerae* populations in an inland location (Dhaka) in cholera-endemic Bangladesh (11, 13).

Here, to understand the role played by subspecies population structure in disease, we compared the *V. cholerae* population from inland Bangladesh with that from the east coast of the USA. This revealed that distribution and abundance of major lineages of *V. cholerae* differed significantly in the two distinct ecosystems. Both globally distributed as well as locally adapted lineages of *V. cholerae* are found in the two environments studied. One of the most striking differences was the presence of several related lineages in Dhaka forming a divergent clade at the base of the *V. cholerae* species in a phylogenomic analysis, which were completely absent in the coastal USA location. Genomic characterization of these lineages reveals that they form a novel species closely related to but distinct from *V. cholerae*. A revision of recent and decades old historical isolates related to this novel species indicates that it has been found in similar environments to pandemic *V. cholerae* for decades and is associated with human infections ranging from septicaemia to choleraic diarrhea.

## Results and Discussion

### Pandemic related strains increase total *V. cholerae* abundance in Dhaka and reduce local diversity

One of the main differences between the *V. cholerae* populations from Oyster Pond (Falmouth, USA) and Dhaka (Bangladesh) is, unsurprisingly, the abundance of the pandemic generating (PG) lineage, which includes strains responsible for the current 7^th^ pandemic. Water samples were previously collected biweekly from seven different sites in the water bodies surrounding Dhaka city for nine continuous months (from June, 2015 to March, 2016), as well as a population from Oyster Pond over the summers of 2008 and 2009 in Cape Cod, Falmouth on the USA east coast (12). Here we compare the *V. cholerae* populations from these two areas to gain insights on the differences between a region that is non-endemic for cholera and experiences strong seasonal variation, with a tropical area endemic for the disease. High-throughput sequencing of *viuB* marker gene amplicons was used to analyse the subspecies composition of *V. cholerae* in these two populations. Amplicons of this gene were annotated following a previously established scheme (11), in which diversity within the *V. cholerae* species is measured based on relative abundance and distribution of *viuB* alleles. Each allele represents a *V. cholerae* lineage, the diversity of which is roughly equivalent to that of a clonal complex as traditionally defined by Multi-locus Sequence Typing (11). A single *viuB* allele (*viuB*-73) can be found to be uniquely associated with the pandemic generating (PG) lineage which is mostly composed of *V. cholerae O1* strains (11). Abundance and distribution of *viuB* alleles in samples collected from the two locations were estimated from *viuB* amplicon sequencing data normalized by quantitative data of *viuB* gene copy numbers determined by qPCR (14). Total abundance of *V. cholerae* in the two locations varied significantly (Kruskal-Wallis test, p<0.1), being almost twice as high on average in Dhaka (2.3×10^5^ gene copies/litre) than in Oyster Pond (1.25 × 10^5^ gene copies/litre) (**Fig. 1A**). However, when PG *V. cholerae* O1 (*viuB-*73) were excluded (quantified independently of other lineages using qPCR of the *rfb*O1 gene), average abundance was very similar in the two locations (Kruskal-Wallis, p<0.01). The PG lineage was the predominant genotype in Dhaka, with an average abundance of 1.4×10^5^ *rfb*O1 gene copies/litre, whereas it was just a minor member of the population in Oyster Pond, with an average abundance of 1.5×10^4^ gene copies/litre (**Fig. 1A**). qPCR analysis confirmed that PG *V. cholerae* O1 present in the Oyster Pond population were non-toxigenic (CTX negative), as opposed to the vast majority of PG *V. cholerae* O1 in Dhaka being toxigenic (CTX positive) (14). Similarity percentage (SIMPER) analysis based on Bray Curtis dissimilarity suggests that the allele most responsible for the overall dissimilarities between Dhaka and Oyster Pond is indeed *viuB*-73. This allele was predominant throughout the nine month sampling period in Dhaka (13), constituting around 60% of the total *V. cholerae* population on average whereas its presence was stochastic in Oyster Pond, with around 5% of the total population (11). Population structure indices (Diversity and Evenness) were significantly lower in Dhaka than in Oyster Pond (Kruskal-Wallis test, P<0.1) (**Fig. 1B**). This indicates a more stable and diverse *V. cholerae* community structure in the coastal location and a less diverse community dominated by fewer alleles in inland Bangladesh (Dhaka), likely because of the dominance of *viuB*-73 in that environment.

**Figure 1:**
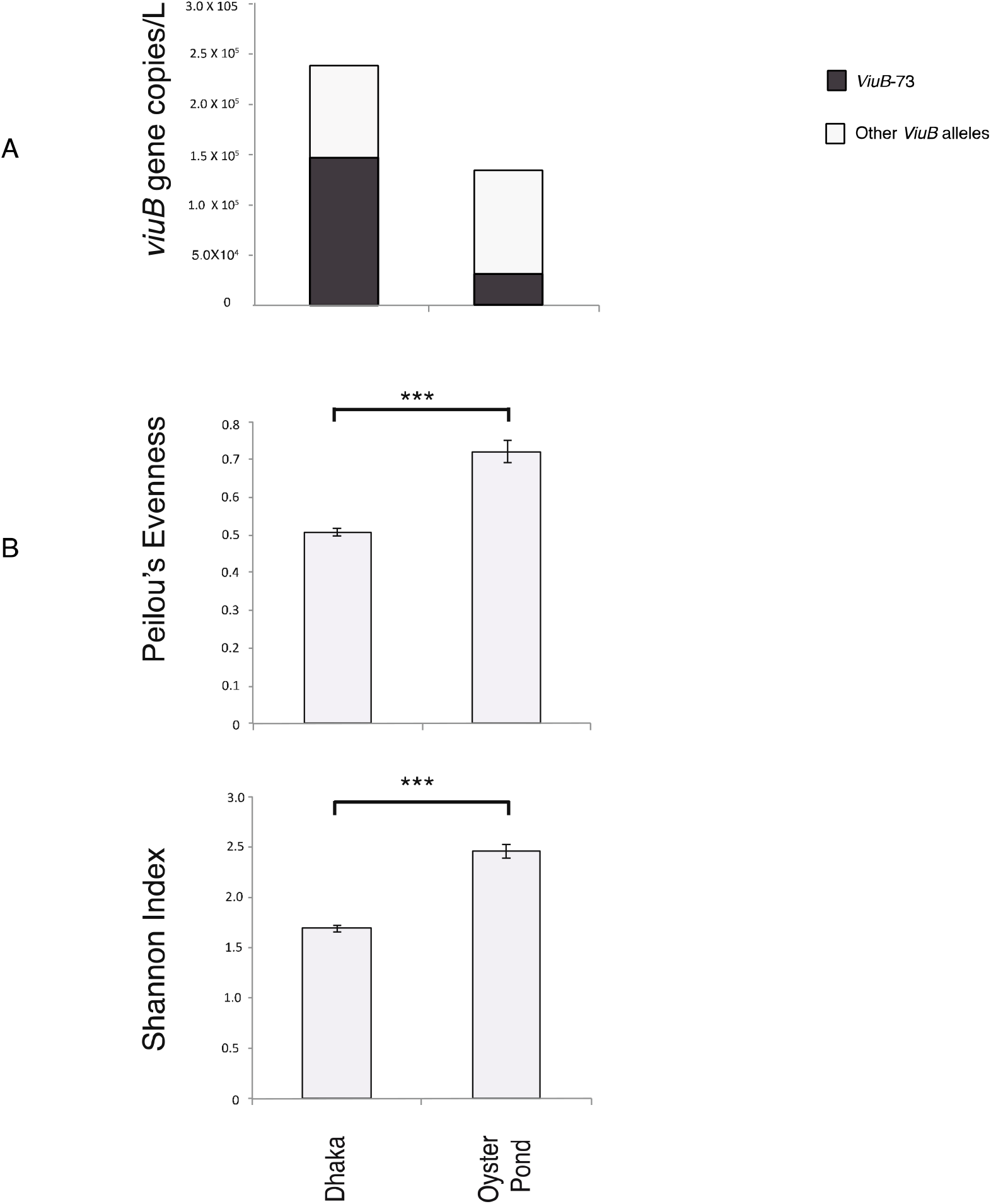
Abundance and Diversity of *Vibrio cholerae* populations in two geographic locations: Dhaka and Oyster Pond. A: Absolute average abundance of *V. cholerae* quantified from qPCR data. Total height of the bar represents total *V. cholerae* (*viuB*), black segment represents *viuB*-73 and clear segment represents other *viuB* alleles. B: Evenness and diversity of the two *V. cholerae* populations measured by Peilou’s evenness and Shannon diversity indices based on analysis of *viuB* alleles. Statistical significance was measured by Kruskal-Wallis test; ***: statistically significant differences (Kruskal-Wallis p<0.1).

Dhaka’s aquatic reservoirs therefore seems to harbour a *V. cholerae* community highly dominated by the PG lineage that is most likely to be affected substantially by human activity. It is one of the most densely populated megacities in the world and has long history of suffering from recurring cholera (15). Sustenance of the cholera causing genotype (PG) in the environment could be the driving factor to shape the overall population of *V. cholerae* in Dhaka. The reduction of intra-species diversity by PG *V. cholerae* in cholera endemic Dhaka could be attributed to the potential selective advantage of colonizing human gut (16), which would result in a constant output to water reservoirs. Type six secretion-mediated killing could also lead to the reduction of diversity, giving advantage to PG *V. cholerae* in a resource limited competitive environment, where PG is a superior competitor to other strains at higher temperatures (17, 18). Environmental conditions, i.e. the lower salinity seen in Dhaka (**Supplementary Table 1**) could also advantage PG strains over others, as they have been shown to be more prevalent in low salt environments relative to other lineages (11).

### A novel divergent lineage is endemic to inland Bangladesh

Besides the PG lineage, the population composition of *V. cholerae* sampled over 6 to 9 months was strikingly different in Dhaka and Oyster Pond. This was determined by using the abundance and distribution data of individual *viuB* alleles from the two locations. Non-metric multi dimensional scaling (NMDS) was performed to compare the two communities and statistical significance of community structure dissimilarity was evaluated using the analysis of similarity (ANOSIM) with a Bray-Curtis distance matrix. In the NMDS plot, samples from Dhaka and Oyster Pond clustered separately and community structure dissimilarity was statistically significant (ANOSIM R=0.75, P value <1%) (**Fig. 2**). Only two major alleles were shared between these locations from a total of 13 *viuB* alleles in Dhaka and 15 alleles in Oyster pond (each individual allele constituting at least 1% of the *V. cholerae* population). The most abundant alleles in Dhaka after *viuB*-73 were *viuB-*06, *viuB-*07, *viuB-*25 and *viuB-*05 (**Fig. 3**). Of these four, three are exclusively found in Dhaka (*viuB-*05, *viuB-*06 and *viuB-*07) and are of particular interest. Together, they composed ∼15% of the average Dhaka *V. cholerae* population and have been found to display higher abundance in sites surrounded by a high human population density and levels of pollution (13).

**Figure 2:**
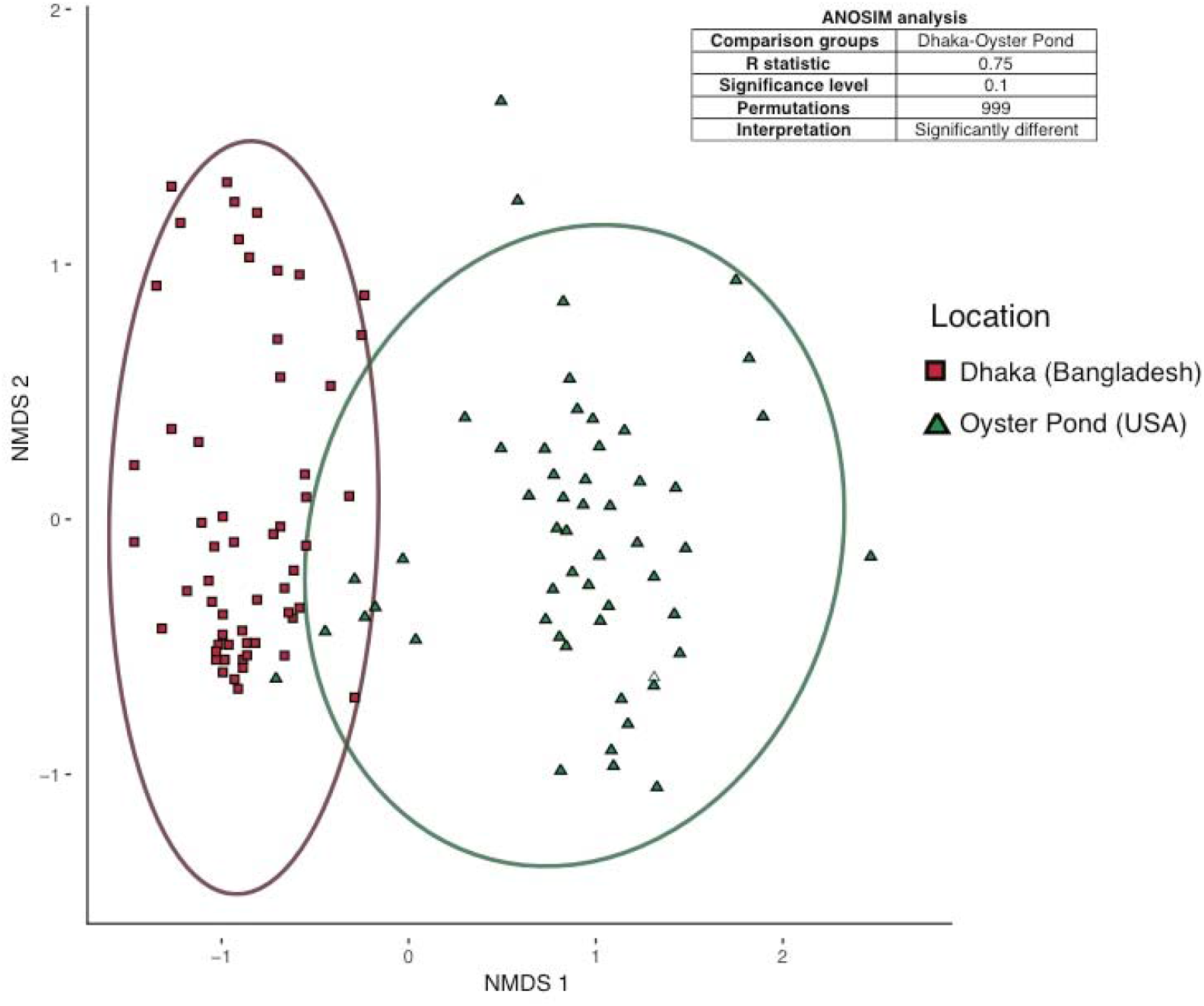
Non-metric multi-dimensional scaling (NMDS) plot comparing beta diversity of *Vibrio cholerae* populations from two aquatic environments. Population compositions were compared using Bray–Curtis dissimilarity matrix with ellipses representing 95% confidence intervals. Dataset was composed of *viuB* gene amplicon sequences normalized by qPCR copy numbers. NMDS plot (stress 0.16) shows distinct clustering of samples from the two locations shown along the first two axes labeled as NMDS1 and NMDS 2. Analyses of similarity (ANOSIM) results are displayed in the box inside the plot describing dissimilarity between pairs of samples from the two locations.

**Figure 3:**
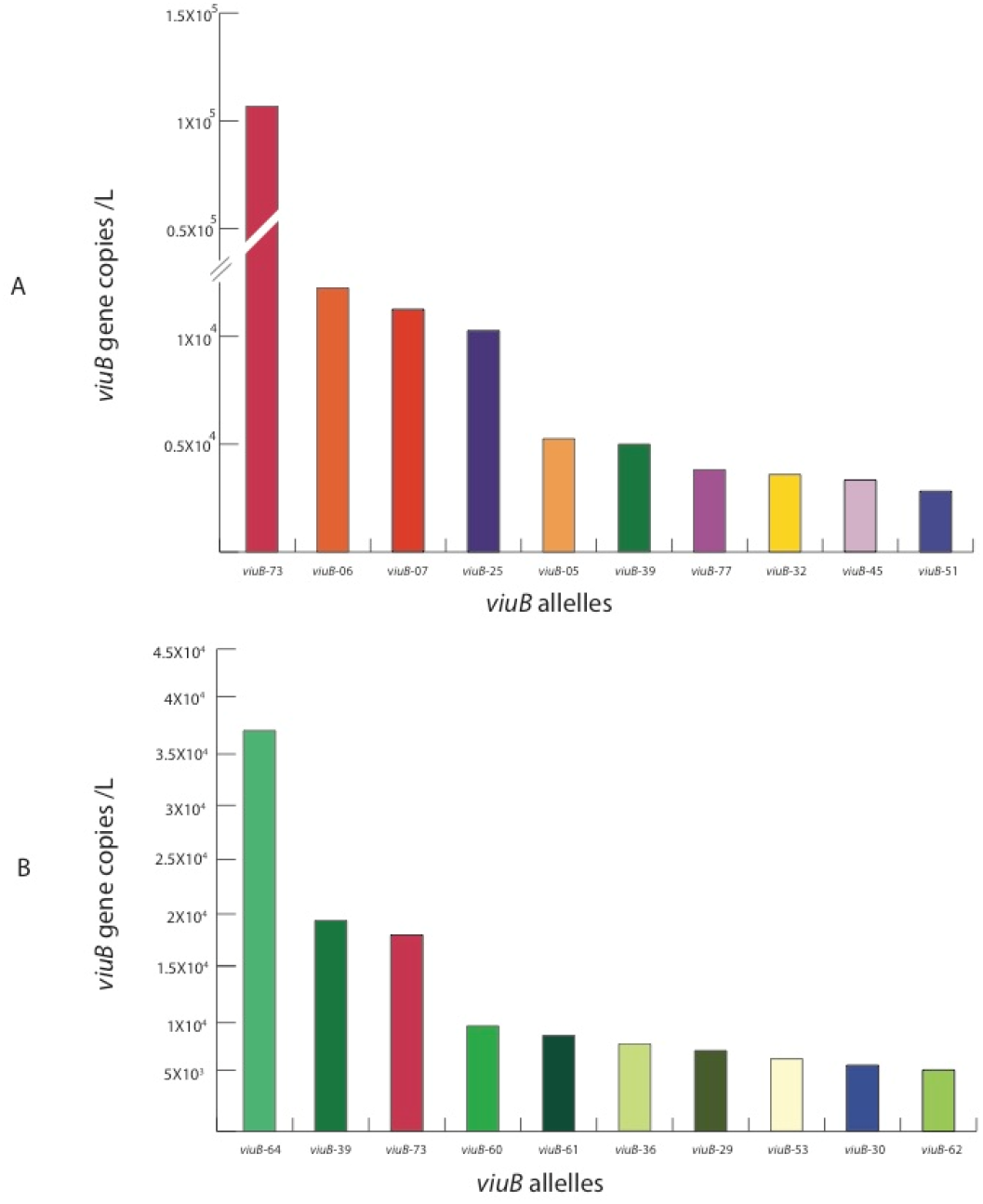
Abundance of the most prevalent *viuB* alleles at two locations: A. Dhaka (Bangladesh); B. Oyster Pond (USA). Total *viuB* gene copy numbers were obtained by qPCR. Relative abundance of each allele was determined by amplicon sequencing. Specific colours were used for individual alleles to be consistent with the scheme described by Kirchberger (11). The ten most abundant alleles for each location were selected for comparison between the two locations.

To have more information on the lineages found in Dhaka, 23 *V. cholerae* strains isolated from the city during the study period were selected for whole genome sequencing: nine *V. cholerae* O1 harbouring the *viuB*-73 allele and fourteen *V. cholerae* non-O1/O139 isolates displaying a diversity of *viuB* alleles. Four strains possessed *viuB* alleles 05, 06, 07 and 08 (EDC690, EDC716, EDC717 and EDC792) and were found to be part of a very long branch occupying a basal position in a global core genome phylogeny compared to the rest of the *V. cholerae* strains (hence termed Long Branch clade or LB) (**Fig. 4**). This phylogenetic group has not been described in any other studies, although other strains from public databases, isolated from different parts of the world, also belong to it. Nine isolates were recovered from human clinical specimens across the United States and reported to the Centre for Disease Control (CDC) as part of the surveillance conducted under the Cholera and Other *Vibrio* Illness Surveillance (COVIS) program (19). Two more isolates originate from stool samples of diarrheal patients in Mozambique in 2008 (20) and one isolate was recovered from a diarrheal patient from Thailand in 1993 and described as *V. cholerae* serogroup O155 (21). Seven additional isolates have been found to belong in the clade for a total of twenty-three as of August 2019 (**Supplementary Table 2**).

**Figure 4:**
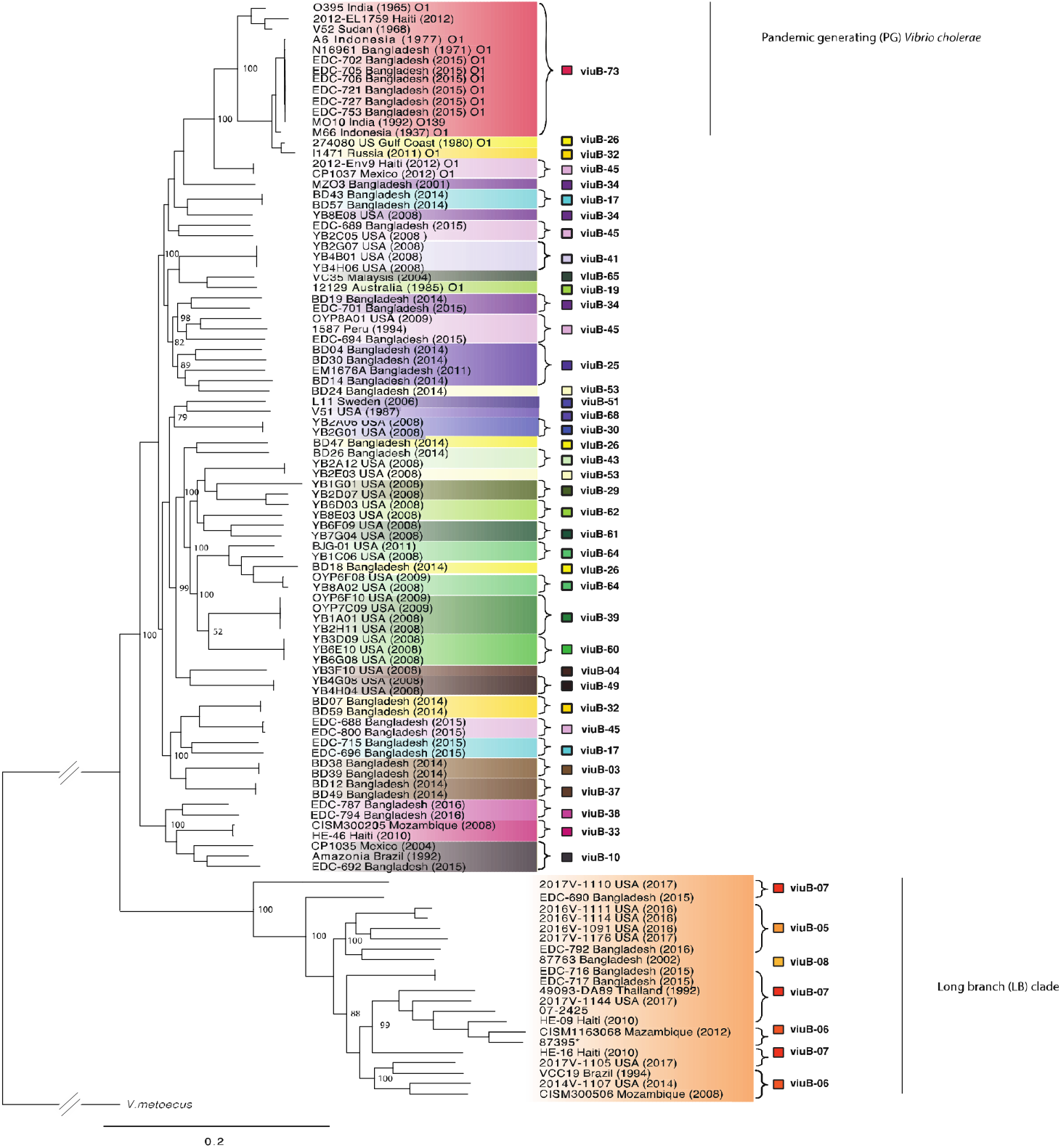
Whole-genome phylogeny of *V. cholerae* strains found in Dhaka and Oyster Pond populations. The phylogenetic tree was inferred using Parsnp v1.2 (46) based on the reference genome of *V. cholerae* O1 El Tor N16961, and includes representative strains from other environments. Leaves of the tree were coloured according to the *viuB* allele found in that particular genome. Statistical support of relevant nodes was estimated by bootstrap analysis (1000 replicates, indicated as a percentage). The scale bar represents nucleotide substitutions per site.

### A sister species to *Vibrio cholerae*?

Comparative genome analysis suggests that LB isolates represent a new species, which would be the closest relative of *V. cholerae* described to date. Based on the genome sequences, G+C content of the strains belonging to LB clade was 46-48.1%, falling within the known range of the genus *Vibrio*. The genomes of twenty-two LB isolates were compared with a set of *V. cholerae* strains containing the same number (n=22) of representatives from both pandemic and non-pandemic lineages (**Supplementary Table 2**). This comparison revealed that genetic distance between LB strains and *V. cholerae* fall below or at the threshold of the species cut-off values. Indeed, Digital DNA-DNA hybridization (dDDH) values ranged from 82-100% within the LB clade and 69-70% with *V. cholerae*, whereas Average Nucleotide Identity (ANI) values ranged from 97-100% within the group and 95-96% with *V. cholerae* strains, respectively (**Supplementary File 1**). DDH values are considered to be the gold standard for species designation and a value of ≤70% presents as an indication that the tested organism belongs to a different species than the type strain(s) used as reference (22). ANI has been proposed as an alternative genomic statistics to DDH and the cut off values of 95-96% has been used for species delineation (23). In this case, all the strains from the LB clade had a dDDH value of 69% and ANI value of 96% when compared with *V. cholerae* type strain N16961. Thus, according to the current species definition (24), the LB clade meets the genotypic criteria to qualify as a candidate for a novel species designation. It also meets the phylogenetic criteria, as it represents a well-defined, well-supported monophyletic clade (**Fig. 4**).

Very recently, genome sequencing efforts of a historical collection of isolates from cholera or cholera-like diseases have identified a strain isolated during the first World War (in 1916) from a soldier convalescent in Egypt as a divergent *V. cholerae* (25). This NCTC30 strain actually belongs to the LB clade found in this study (**Fig. 5**). Interestingly, NCTC30 was initially designated as “*Vibrio paracholerae*” and the disease caused was described as choleraic and termed as ‘paracholera’ (26). To honour its history, we propose the name *Vibrio paracholerae* sp. nov. (EDC-792^T^) for this novel species..

**Figure 5:**
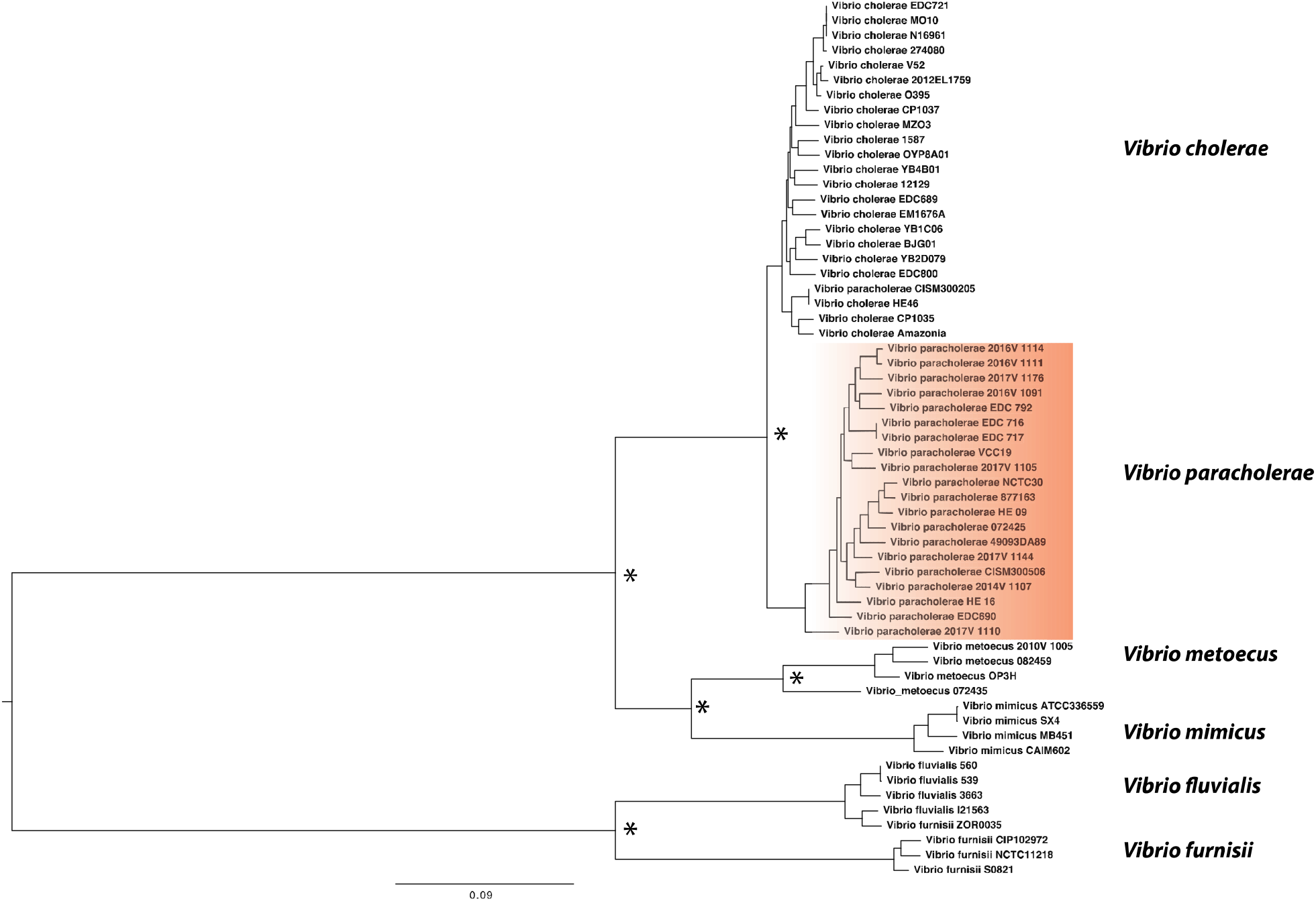
Whole-genome phylogenetic tree of *V. paracholerae* along with its closest sister species. The maximum likelihood phylogenetic tree was constructed from the core genome alignment of ≈2.1M bp using GTR gamma substitution model. Corresponding nodes with relevant Bootstrap support over 70% from the 100 replicates were indicated with *. The scale bar represents nucleotide substitutions per site.

To confirm that *V. paracholerae* sp. nov. was indeed a novel species, its phenotypic traits were compared to those from the most closely related species: *V. cholerae* and *V. metoecus*. Their ability to catabolise 190 different carbon sources and response to 96 chemicals and antimicrobials were determined using Biolog Phenotypic Microarray (PM) plates. Four *V. paracholerae* strains were examined, two of environmental origin in Bangladesh (EDC 690, EDC792) and two from clinical sources in the USA (2016V-1114, 2016V-1091). These were compared with four *V. cholerae* (N16961, V52, YB3B05, YB8E08) and four *V. metoecus* strains (082459, OP6B, OP4B, OP3H). Although the *V. paracholerae* sp. nov. strains resembled *V. cholerae* in most biochemical and growth characteristics, they clearly differed for some phenotypic characteristics (**Table 1**). All four *V. paracholerae* sp. nov. strains tested could utilize α**-**cyclodextrin as a sole carbon source, whereas none of the tested *V. cholerae* strains could. Cyclodextrin utilization requires a specific category of amylases, which has not been reported in *V. cholerae* so far (27). *In silico* analysis revealed that *V. paracholerae* strains possess a gene cluster (genes 03367 to 03379 in the NCTC30 genome, NZ_LS997867) containing homologs of genes encoding cyclomaltodextrin glucanotransferase (*amyM*), ABC transporter MalK (*malK*), glycosidase MalE (*malE*), glucosamine N-acetyltransferase, cyclodextin specific porin (*cycA*), cyclodextrin binding protein (*cycB*), cyclodextrin transport system permease (*malF*), cyclodextrin transport system permease (*malD/malG*) and neopullulanase (*nplT*) (**Table 2**). Only two (9%) *V. cholerae* strains in our dataset (n=22) and a similar percentage in the NCBI database possessed this cluster whereas 100% of the *V. paracholerae* sp. nov. strains (n=22) harboured it. This cluster might be associated with the cyclodextrin degradation phenotype, as reported previously (28). In contrast with *V. cholerae*, 75% (3 out of 4) of *V. paracholerae* sp. nov. strains tested were found to be lacking the ability to utilize D-mannose, L-aspartic acid, citric acid, alpha keto glutaric acid and mono-methyl succinate (**Table 1**). D-mannose was found to be readily utilized by both *V. cholerae* and *V. metoecus* tested in this study and previous literature reported that ∼80% of *V. cholerae* are capable of utilizing this sugar (3). The gene cluster encompassing *manP* to *manA* (VC1820 to VC1827 in N16961 genome, AE008352.1), including the well-known mannose-6 phosphate isomerase (*manA*) gene required for this process (29), was present in all the tested *V. cholerae* (n=22) and *V. metoecus* (n=4) strains, whereas it was found in only ∼40% (9/22) of *V. paracholerae* sp. nov. strains **(Table 2**). We could not find the genetic basis for the other phenotypic differences between *V. paracholerae* sp. nov. and *V. cholerae. V. paracholerae* is similar to *V. cholerae* in N-Acetyl-D-Galactosamine and D-glucuronic acid utilization tests and acetoin production, which differentiates both species from *V. metoecus* (7). Resistance to 96 drugs or metals were also tested at different concentrations, and *V. cholerae* and *V. paracholerae* sp. nov. showed similar profiles in most, although three chemicals elicited differential responses by the two species. *V. paracholerae* sp. nov. strains were resistant to cadmium chloride, sodium selenite and dichlofluanid in contrast to the sensitivity of the *V. cholerae* strains towards those chemicals (**Supplementary Table 3**). These differences in carbon source utilization capability and response to antimicrobial chemicals could be crucial in defining the ecological preferences of *V. paracholerae* sp. nov. and interactions with its more famous sister species.

**Table 1:** Phenotypic traits differentiating *Vibrio paracholerae* sp. nov. from its closest relatives *Vibrio cholerae* and *Vibrio metoecus*. Strains: 1, EDC 792; 2, EDC 690; 3, 2016V-1111; 4, 2016V-1091; 5, N16961; 6, V52; 7, YB3B05; 8, YB8E08; 9, Vm 082459; 10, OP6B; 11, OP4B; 12, OP3H. +, Growth/positive test result; -, no growth/negative test result; ND, not determined. †Results for *V. cholerae* and *V. metoecus* strains were obtained from Kirchberger *et al* (7). †Results for *V. cholerae* and *V. metoecus* strains were obtained from Kirchberger (7)

| Phenotypic test | *Vibrio paracholerae* sp. nov. |  |  |  | *Vibrio cholerae* |  |  |  | *Vibrio metoecus* |  |  |  |
| --- | --- | --- | --- | --- | --- | --- | --- | --- | --- | --- | --- | --- |
|  | 1 | 2 | 3 | 4 | 5 | 6 | 7 | 8 | 9 | 10 | 11 | 12 |
| α-cyclodextrin | + | + | + | + | - | - | - | - | + | + | - | + |
| Pectin | + | + | + | + | - | - | - | - | + | + | + | + |
| Mono methyl succinate | - | - | + | - | + | + | + | + | - | - | - | - |
| D-Mannose | - | - | + | - | + | + | + | + | + | + | + | + |
| L-aspartic acid | - | - | + | - | + | + | + | + | + | + | + | + |
| Citric acid | - | - | + | - | + | + | + | + | + | + | + | + |
| α-keto glutaric acid | - | - | + | + | + | + | + | + | - | - | + | - |
| N-acetyl-D-galactosamine | - | - | - | - | - | - | - | - | + | + | + | + |
| D-glucuronic acid | - | - | - | - | - | - | - | - | + | + | + | + |
| Acetoin production | + | + | + | + | + | + | + | + | - | - | - | - |

**Table 2:**
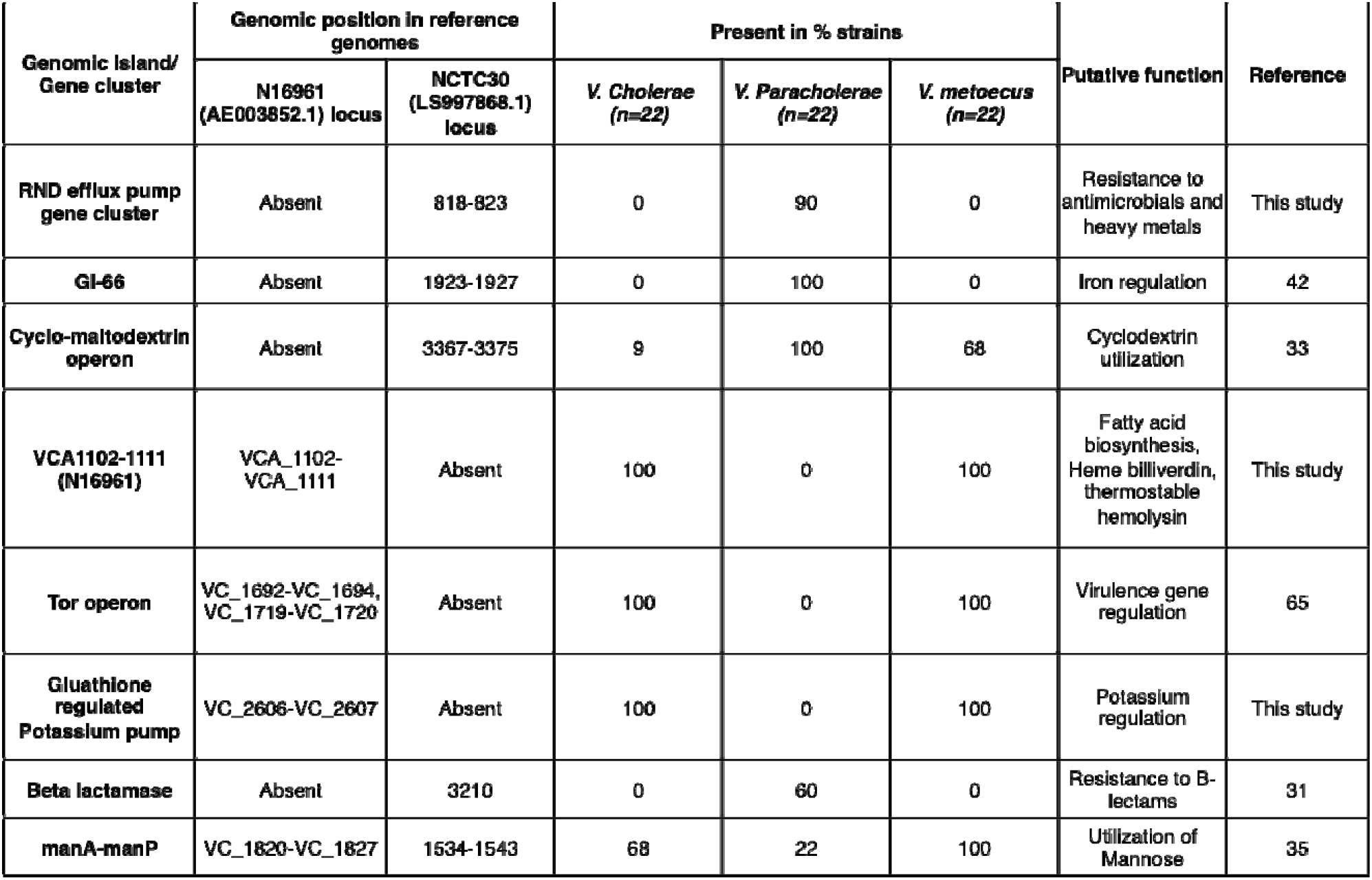
Major genetic traits differentiating *Vibrio paracholerae* sp. nov. from its closest relatives: *Vibrio cholerae* and *Vibrio metoecus*. VC, *Vibrio cholerae*; VP, *Vibrio paracholerae* sp. nov.; VM, *Vibrio metoecus*. Reference genomes N16961 (V. cholerae) and NCTC30 (*V. paracholerae* sp. nov.) were used for determining locus positions of the gene clusters.

### A potential threat to humans?

To be a successful disease-causing agent to humans, a bacterial pathogen of aquatic origin needs to have the ability to survive in the environment and colonize the human body. In cholera endemic Dhaka, *V. paracholerae* sp. nov. has been found to exist abundantly in local water reservoirs. In one particular site, the number even surpassed that of PG *V. cholerae*, which was otherwise the most predominant lineage found in Dhaka (13). That site (Kamrangir char) happened to be the most densely populated region among those sampled, indicating a possible link between human population and the prevalence of *V. paracholerae* sp. nov. This raises the possibility of adaptation to the human gut as an alternate niche and an important factor in its ecology. Association of the members of the species with cholera-like cases, such as in the case of the historical strain NCTC30, and isolation from clinical/human stool samples from different parts of the world would suggest their pathogenic potential to humans. To assess this potential, *V. paracholerae* sp. nov. strains were screened for the presence of known virulence-related genes and islands often found in *V. cholerae* **(Supplementary Table 4)**. *V. paracholerae* sp. nov. strains lack CTX, VPI1 and VPI2; three major elements known to be essential for *V. cholerae* to cause cholera (2). They also lack a cluster of genes (VC1692, VC1694, VC1719 and VC1720 in the N16961 genome) encoding proteins for the ‘Tor operon’ required for trimethylamine N-oxide respiration in *V. cholerae*. The genes in this operon have been shown to be crucial for cholera toxin production, cytotoxicity and intestinal colonization of *V. cholerae* in infant mouse model (30). The Tor operon was found in 100% of *V. cholerae* (n=22) and *V. metoecus* (n=4) strains in our dataset, which indicate that it was likely lost in the *V. paracholerae* sp. nov. phylogenetic branch, possibly impacting their interaction with eukaryotes and distinguishing it from its sister species.

All the *V. paracholerae* sp. nov. strains in our dataset possessed the RTX toxin gene cluster, a virulence factor for *V. cholerae* known to have a role in interaction with eukaryotes (31). Interestingly, five *V. paracholerae* sp. nov. strains (22%) (including NCTC30) possess Type Three Secretion System genes, an established virulence factor for non-pandemic *V. cholerae* (32). Six strains possessed an SXT element, which is found in most PG *V. cholerae* strains since 2001 and is believed to be involved in improved fitness in 7^th^ pandemic El Tor *V. cholerae* (9, 33). Additionally, 50% of the *V. paracholerae* sp. nov. strains also contained genes for the recently discovered cholix toxin, thought to be an important virulence factor for *V. cholerae* (34).

Apart from the known virulence genes usually found in *V. cholerae*, gene content analysis revealed a few species-specific genetic traits in *V. paracholerae* sp. nov. Two genes were present in all 22 *V. paracholerae* sp. nov. strains, with no homolog found in any *V. cholerae* strains. These two genes encode a lysR family transcriptional regulator (WP_001924807.1) and HAD_IB family hydrolase (WP_071179638.1). Both of these genes are part of a previously reported genomic island (GI-66) found in *V. albensis* (35). This GI contains iron-related regulatory genes that can be significant in regulation of iron scavenging in this group of organisms. Iron acquisition is thought to be an important aspect for regulation of virulence as well as host selectivity/ specificity (36). Most (90%) of *V. paracholerae* sp. nov. strains also harboured a novel RND efflux pump (**Table 2**), thought to be critical for intrinsic and induced antimicrobial resistance, virulence gene expression, colonization in animal host and environmental regulation of stress response (37). Efflux pumps have been proposed to be important for expelling bile out of the cell, and the resulting bile resistance would be key to overcoming this challenge inside the human gut (38). RND efflux pumps have specifically been found to confer increased bile resistance in other gram negative bacteria (39). The novel RND gene cluster is absent in both *V. cholerae* and *V. metoecus* but homologs have been found in the halophilic bacteria *V. cincinnatiensis* (40) and a bile associated isolate of *V. fluvialis* (41). Other than efflux pumps, ToxR and TolC have been proposed to be crucial for bile resistance, and like *V. cholerae* strains, all the *V. paracholerae* sp. nov. strains possess both genes. All these factors make *V. paracholerae* sp. nov. a potential candidate for a species adapted to the human gut and underscores the importance of studying their biology in greater detail.

### Interaction of *Vibrio paracholerae* sp. nov. with pandemic *Vibrio cholerae* impacts the ecology and evolution of both species

Horizontal gene transfer (HGT) among species sharing an ecological niche can have a major impact on their evolution (42). As *V. paracholerae* sp. nov. (VP) co-exists with *V. cholerae* (VC) in natural ecosystems (at least in Dhaka), it is expected that HGT could take place between these two groups. To assess the propensity of interspecies HGT, potential gene transfer events within two groups (VC and VP) were inferred based on phylogenetic congruence of individual genes. Maximum likelihood (ML) trees were constructed for each of the core and accessory gene families present in at least two strains from each group. A gene transfer was hypothesized if a member of a group clustered with members of the other group in a clade, and the gene tree could not be partitioned into perfect clades, which must consist of all members from the same group and only of that group (8, 43). In our groups of 22 VC and 22 VP strains, 216 HGT events were hypothesized involving 82 gene families from VC to VP, but only 62 events from VP to VC involving 33 gene families. All of the core genes transferred from VP to VC were acquired by strains outside of the PG group. In the case of accessory genes, we could infer 82 potential transfer events from VC to VP and 54 events from VP to VC. Only 4 events involved strains belonging to the PG clade. Thus, gene transfer directionality was biased from VC to VP, VP being the recipient of HGT in most cases. Lower rate of HGT towards *V. cholerae* was previously reported in case of the co-occurring *V. metoecus*, which has a lower abundance in the environment (8). This gene transfer bias could be attributed to the dominance of *V. cholerae* in cholera endemic region, as it is generally more abundant than *V. paracholerae* sp. nov. and therefore more likely to be a DNA donor (14). Among the accessory genes transferred from *V. cholerae* to *V. paracholerae* sp. nov., there were proteins related to O antigen synthesis, T6SS, iron regulation, chaperone and multi-drug resistance and putative metabolic functions. There are examples of a single gene or even a small set of nucleotides within a gene acquired via HGT impacting the ecology and pathogenicity of bacterial lineages (42, 44). Thus, the HGT events in *V. paracholerae* sp. nov. underscore the possibility for species co-existing with PG *V. cholerae* to acquire virulence and fitness-related genes to become pathogenic to human and/or novel ecological traits. Gene transfer events have led to the rise of virulent *V. cholerae* before, a great example being the rise of *V. cholerae* O139. The latter emerged in Bangladesh and India in 1992 and is thought to have originated via genetic recombination of O-antigen region from a serogroup O22 strain to a serogroup O1 El Tor strain (45). After its emergence, *V. cholerae* O139 remained an important cause of widespread cholera epidemics in that region until 2004, along with *V. cholerae* O1 El Tor (45). Interestingly, it appears that among the accessory genes inferred as subject to HGT from VP to VC, genes encoding UDP-glucose 4-epimerase (EC 5.1.3.2) and UDP-N-acetylgalactosaminyltransferase could be result of transfers in an ancestor of O139 strain MO10 from the *V. paracholerae* sp. nov. clade (**Fig. 6**). Both genes are involved in O-antigen biosynthesis and could be of significance in the emergence and evolution of *V. cholerae* O139 as a human pathogen and pandemic agent. Even though it will require further investigation to find out how and to what extent *V. paracholerae* sp. nov. as a species contributed to the emergence and evolution of *V. cholerae* O139, these transfer events could be considered as examples of how interaction of this close relative with *V. cholerae* could impact the epidemiology of cholera.

**Figure 6:**
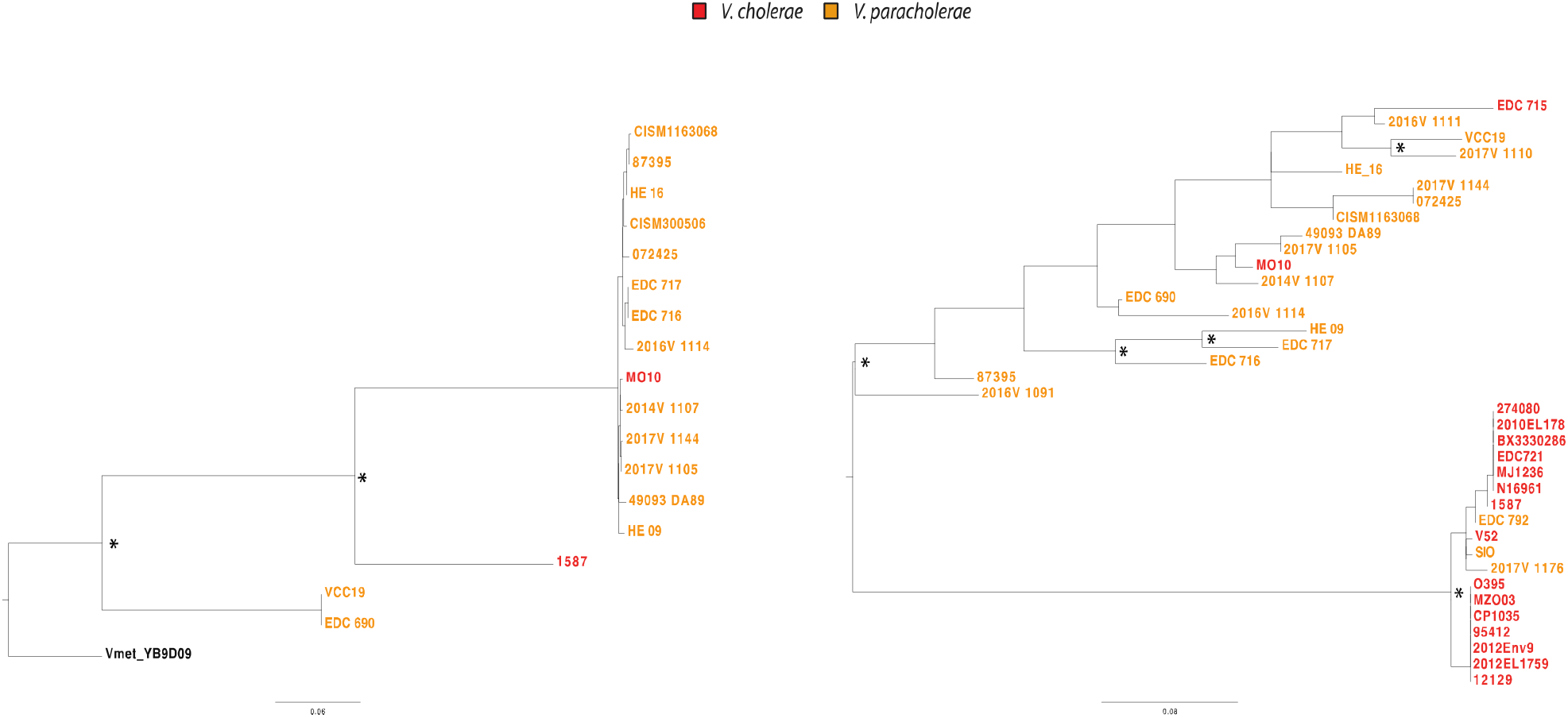
Phylogenetic tree of O-antigen cluster genes found in *V. paracholerae* and *V. cholerae*. Maximum likelihood trees were constructed using A) 705 bp nucleotide alignment of the gene encoding UDP-glucose 4-epimerase and B) 564 bp alignment of the gene encoding UDP-N acetylgalactosaminyltransferase. Nodes with relevant Bootstrap support over 70 of 100 replicates are indicated with *. The scale bar represents nucleotide substitutions per site.

## Conclusions

Culture-independent analysis below the species level in inland cholera endemic and coastal non-endemic locations in distinct geographic settings identified differences in the population structures present in these environments. It revealed that human influences are likely to be a major factor shaping communities of that species in cholera endemic areas. In urban tropical Dhaka, found in inland Bangladesh, PG *V. cholerae* was abundant and continuously present, but accompanied by members of a related but phylogenetically distinct clade, which could represent a novel species. The abundance of this putative species, ‘*Vibrio paracholerae* sp. nov.*’*, in Dhaka and its absence from Oyster Pond on the USA East Coast, indicates that it is not a ubiquitous member of aquatic communities. In addition to those identified here from the COVIS program in the USA and from Mozambique and Thailand, several strains of *Vibrio* spp. have been very recently isolated from clinical cases in China and Korea which would belong to this species according to the genome sequence similarities they share with strains analyzed here (**Supplementary Table 5**). An indirect association of their abundance with human population density indicates that they could be adapted to the human gut in cholera endemic areas (13). They could therefore occasionally become pathogenic by acquiring pathogenicity gene clusters or cause opportunistic infections in vulnerable individuals. The history, biology, genetic traits and coexistence with a pathogenic sister species makes it a risk as an emerging human pathogen. Its potential contribution to the evolution of new pathogenic variants of *V. cholerae* (such as PG lineage O139) and likely influence on their population structure highlights the importance of studying this novel species in the context of a globally distributed infectious disease.

## Data availability

The sequences reported in this article have been deposited in the NCBI database under bioproject number: PRJNA598367.

## Acknowledgement

This work was supported by the Natural Sciences and Engineering Research Council (NSERC) of Canada (to YB); the Integrated Microbial Biodiversity program of the Canadian Institute for Advanced Research (to YB); federal appropriations to the Centers for Disease Control and Prevention through the Advanced Molecular Detection Initiative (to CLT); and graduate student scholarships from Alberta Innovates – Technology Futures (to MTI), the NSERC Canada Graduate Scholarship – Doctoral Program (to TN), and the Bank of Montréal Financial Group (to FDO). The funders had no role in study design, data collection and interpretation, or the decision to submit the work for publication. The findings and conclusions in this report are those of the authors and do not necessarily represent the official position of the Centers for Disease Control and Prevention. MA of icddr, b thanks the government of Bangladesh, Canada, Sweden and United Kingdom for providing unrestricted core support.

## Materials and Methods

### Sample collection and processing

Environmental water samples were collected every two weeks between June 2015 and March 2016 from seven points along the water bodies surrounding Dhaka city, which is located in the central part of Bangladesh (23.8103° N, 90.4125° E). One-time water samples were collected from two natural coastal water bodies in Mathbaria (22.2920° N, 89.9580° E) and Kuakata (21.8210° N, 90.1214° E), which are geographically adjacent to the coast of the Bay of Bengal and approximately 200 km and 250 km southwest of Dhaka, respectively. One liter of water was collected from each sites in sterile Nalgene bottles placed in an insulated plastic box, and transported at ambient air temperature from the site of collection to the central laboratory of the International Center for Diarrheal Disease Research, Bangladesh (ICDDR,B), in Dhaka. Oyster pond sampling was performed at the same spots and approximates same time of the day in the pond and the nearby lagoon connected to the ocean in monthly intervals from June to October as described by Kirchberger (12). 50 liters of water were filtered through 0.22μm sterivex filters (Mo Bio Laboratories Inc., Carlsbad, CA, USA) for the collection of biomasses. Genomic DNA was extracted from the biomass using the protocol described by Wright (47).

### Isolation and identification of isolates

Bacterial isolates were recovered as described elsewhere (48). Briefly, water samples were enriched in APW (Difco Laboratories, Detroit, Mich.) at 37°C for 6 to 8 h before plating. About 5 μl of enriched APW broth was streaked, using an inoculating loop, onto both thiosulfate-citrate-bile-salts-sucrose (TCBS) and TTGA and incubated at 37°C for 18 to 24 h. Colonies with the characteristic appearance of *V. cholerae* were confirmed by standard biochemical and serological tests (and, in the case of the latter, by testing with polyvalent and monoclonal antibodies specific for *V. cholerae* O1 or O139) and, finally, by PCR.

### Phenotypic tests

For the comparison of phenotypic characteristics, Biolog phenotypic microarray plates PM1, PM2A, PM14A, PM16A, PM18C were used (49). Overnight cultured bacterial colonies were inoculated into Biolog IF-0a Base medium to reach 85 % turbidity followed by 1:200 dilution aliquoted into IF-10b medium supplemented with Dye Mix A as indicated by the manufacturer instructions. The mixture was then added into wells of Biolog PM1 and PM2A plate containing various carbon sources and PM14A, PM16A and PM18C plates containing substrates of various antimicrobials and heavy metal salts. The incubation and monitoring of the growth of inocula were done for 96 h in the presence of sole carbon source or the heavy metals, growth causes reduction of the dye, resulting in purple colour formation.

### Quantitative PCR (qPCR)

Estimation of *Vibrio cholerae* number was done using qPCR following the protocol described elsewhere (14). Briefly, Target probe for *viuB*, 5’-/56-FAM/TCATTTGGC/ZEN/CAGAGCATAAACCGGT/ 3IABkFQ/-3’, forward primer 5’-TCGGTATTGTCTAACGGTAT-3’, and reverse primer 5’-CGATTCGTGAGGGTGATA-3’ was used. The volume of the PCR reaction was 10 μl containing 5 μl of 2× Dynamite qPCR master mix (MBSU, University of Alberta, Edmonton, Canada), 1 μl of each of 500 nM primer-250 nM probe mix, 1 μl of molecular grade water and 2 μl of DNA template. Real-time quantitative PCR was performed under the following conditions: initial primer activation at 95 °C for 2 min followed by 40 cycles of 95 °C for 15 s, 60 °C for 1 min in Illumina Eco Real-Time PCR system.

### Amplicon sequencing

Amplicon sequencing of *viuB* gene was performed following the method described elsewhere (11). To amplify 293 bp of the *viuB* region from DNA extracted from water samples, a touchdown PCR was performed using 0.5 μL each of 10 pmol forward and reverse primers (for *viuB*: *viuB*2f 5’-CCGTTAGACAATACCGAGCAC-3’ and *viuB*5r 5’-TTAGGATCGCGCACTAACCAC-3’), 0.4 μL of 10 mM dNTP mix (ThermoFisher), 0.4 of μL Phire Hot Start II DNA Polymerase (ThermoFisher), 0.5 μL of molecular biology grade bovine serum albumin (20 mg/mL, New England Biolabs), 5 μL of 5× Phire Buffer, and 2 μL of template DNA. The PCR reaction was performed as follows: initial denaturation at 98°C for 4 min; followed by 10 cycles of denaturation at 98°C for 10 sec, annealing at 60°C for 6 sec (reduced by 1°C per cycle), and extension 72°C for 1 sec; followed by 23 cycles of denaturation at 98°C for 10 sec, annealing at 50°C for 6 sec (reduced by 1°C per cycle), and extension at 72°C for 1 sec; and a final extension at 72°C for 1 min. In preparation for sequencing, dual-indexed sequences were tagged using indices developed by Kozich (50) as follows: 2 μL of preceding *viuB* PCR amplification reaction were used as template for a tagging PCR reaction; initial denaturation at 98°C for 30 sec; followed by two cycles of denaturation at 98°C for 10 sec, annealing at 55°C for 6 sec, and extension at 72°C for 1 sec; and final extension at 72°C for 1 min. Eight tagging reactions were performed for each sample and products were pooled and ran on a 2% agarose gel in 1× Tris-Acetate-EDTA buffer. The appropriate bands (428 bp) were cut out of the gel. PCR products were then purified using Wizard SV Gel and PCR Clean-Up System (Promega) according to the instructions by the manufacturer. Concentration of clean PCR products was then measured using a Qubit Fluorometer (ThermoFisher) with a Qubit dsDNA HS Assay Kit (ThermoFisher) and pooled together in equal concentrations (>10 ng/μL). The pooled samples were then concentrated using a Wizard SV Gel and PCR Clean-Up System (Promega). Quality control of the pooled and concentrated sample was done using an Agilent 2100 Bioanalyzer. Sequencing was performed using Illumina MiSeq technology with a v3 (600 cycles) reagent kit.

### Amplicon sequence analysis

De-multiplexed raw reads from the sequencing run were processed in R (51) using the DADA2 pipeline 1.4.0 (52). First 10 bp of forward and reverse reads were trimmed and reads with a maximum expected error rate >1 was discarded. Chimera detection implemented in DADA2 was then performed on pooled samples. To account for the possibility of real chimeras between protein coding genes from closely related organisms (due to recombination or homoplasic mutations), chimeras were compared with a reference dataset of *viuB* alleles found in 782 sequenced *V. cholerae* genomes (obtained from GenBank). Only *viuB* alleles composed of more than 1,000 reads, found in multiple samples (with an average of 100,000 reads per sample) were considered for further analysis. Samples were rarefied to the level of the sample with the lowest reads using mothur 1.39.5 (53), and further analysis was performed in R, with statistical tests and distance calculations performed using the VEGAN 2.4-6 package (54). Bray-Curtis similarity was calculated based on relative read abundance of each allele in different samples in Primer-E Software Suite and used for similarity percentage (SIMPER) and non-metric multi dimensional scaling (NMDS) analysis.

### Whole-genome sequencing and core genome phylogeny

The genomes of 23 strains from Dhaka belonging to various *viuB* genotypes were chosen for whole-genome sequencing as described by Orata (8). Sequencing libraries were prepared from the genomic DNA using the Nextera XT DNA library preparation kit (Illumina, San Diego, CA, USA) and sequenced using Illumina MiSeq sequencing platforms (2□×□250-bp paired-end reads). Quality control and *de novo* assembly of the reads were done using default parameters in CLC Genomics workbench 7 (Qiagen). Whole-genome alignment was performed using Mugsy v1.2.3 (55) with default parameters, and a maximum likelihood tree was built from this alignment using RaxML v8 (56) under the GTR+GAMMA model with 100 bootstrap replicates. Additional *V. cholerae* genomes were downloaded from GenBank. The maximum likelihood phylogenomic tree was constructed from the alignment of locally collinear blocks (2,094,734 bp) using GTR gamma substitution model with 100 bootstrap replicates.

### Comparative genomic analysis

The genome sequences were annotated with RAST 2.0 (57). Genomic distances were calculated in Geneious (58). Core and accessory genes were determined with BPGA finding orthologous protein-coding genes clustered into families based on a 30% amino acid sequence identity (59). Group specific genes were clustered using a custom-made Python program.

## References

1. Ali M, Lopez AL, Ae You Y, Eun Kim Y, Sah B, Maskery B, Clemens J. 2012. The global burden of cholera. Bull World Health Organ https://doi.org/10.2471/blt.11.093427.

2. Kaper JB, Morris Jr. JG, Levine MM. 1995. Cholera. Clin Microbiol Rev 8:48–86.

3. Davis BR, Fanning GR, Madden JM, Steigerwalt AG, Bradford HB, Smith HL, Brenner DJ. 1981. Characterization of biochemically atypical Vibrio cholerae strains and designation of a new pathogenic species, Vibrio mimicus. J Clin Microbiol https://doi.org/10.1128/jcm.14.6.631-639.1981.

4. Hasan NA, Grim CJ, Haley BJ, Chun J, Alam M, Taviani E, Hoq M, Munk AC, Saunders E, Brettin TS, Bruce DC, Challacombe JF, Detter JC, Han CS, Xie G, Nair GB, Huq A, Colwell RR. 2010. Comparative genomics of clinical and environmental Vibrio mimicus. Proc Natl Acad Sci U S A https://doi.org/10.1073/pnas.1013825107.

5. Neogi SB, Chowdhury N, Awasthi SP, Asakura M, Okuno K, Mahmud ZH, Islam MS, Hinenoya A, Nair GB, Yamasaki S. 2019. Novel cholera toxin variant and ToxT regulon in environmental Vibrio mimicus isolates: Potential resources for the evolution of Vibrio cholerae hybrid strains. Appl Environ Microbiol https://doi.org/10.1128/AEM.01977-18.

6. Haley BJ, Grim CJ, Hasan NA, Choi SY, Chun J, Brettin TS, Bruce DC, Challacombe JF, Detter JC, Han CS, Huq A, Colwell RR. 2010. Comparative genomic analysis reveals evidence of two novel Vibrio species closely related to V. cholerae. BMC Microbiol https://doi.org/10.1186/1471-2180-10-154.

7. Kirchberger PC, Turnsek M, Hunt DE, Haley BJ, Colwell RR, Polz MF, Tarr CL, Boucher Y. 2014. Vibrio metoecus sp. nov., a close relative of Vibrio cholerae isolated from coastal brackish ponds and clinical specimens. Int J Syst Evol Microbiol 64:3208–3214.

8. Orata FD, Kirchberger PC, Méheust R, Jed Barlow E, Tarr CL, Boucher Y. 2015. The dynamics of genetic interactions between Vibrio metoecus and Vibrio cholerae, two close relatives co-occurring in the environment. Genome Biol Evol https://doi.org/10.1093/gbe/evv193.

9. Chun J, Grim CJ, Hasan NA, Lee JH, Choi SY, Haley BJ, Taviani E, Jeon YS, Kim DW, Lee JH, Brettin TS, Bruce DC, Challacombe JF, Detter JC, Han CS, Munk AC, Chertkov O, Meincke L, Saunders E, Walters RA, Huq A, Nair GB, Colwell RR. 2009. Comparative genomics reveals mechanism for short-term and long-term clonal transitions in pandemic Vibrio cholerae. Proc Natl Acad Sci U S A 106:15442–15447.

10. Islam MT, Alam M, Boucher Y. 2017. Emergence, ecology and dispersal of the pandemic generating Vibrio cholerae lineage. Int Microbiol 20:106–115.

11. Kirchberger PC, Orata FD, Nasreen T, Kauffman KM, Tarr CL, Case RJ, Polz MF, Boucher YF. 2020. Culture-independent tracking of Vibrio cholerae lineages reveals complex spatiotemporal dynamics in a natural population. Env Microbiol https://doi.org/10.1111/1462-2920.14921.

12. Kirchberger PC, Orata FD, Barlow EJ, Kauffman KM, Case RJ, Polz MF, Boucher Y. 2016. A Small Number of Phylogenetically Distinct Clonal Complexes Dominate a Coastal Vibrio cholerae Population. Appl Env Microbiol 82:5576–5586.

13. Tania Nasreen, Mohammad Tarequl Islam, Kevin Y. H. Liang, Fatema-Tuz Johura, Paul C. Kirchberger, Marzia Sultana Rebecca J. Case MA and YFB. Environmental factors influence subspecies population structure of Vibrio cholerae in Dhaka, Bangladesh. Unpublished.

14. Nasreen T, Hussain NAS, Islam MT, Orata FD, Kirchberger PC, Case RJ, Alam M, Yanow SK, Boucher YF. 2020. Simultaneous quantification of vibrio metoecus and vibrio cholerae with its o1 serogroup and toxigenic subpopulations in environmental reservoirs. Pathogens https://doi.org/10.3390/pathogens9121053.

15. Rafique R, Rashid MU, Monira S, Rahman Z, Mahmud MT, Mustafiz M, Saif-Ur-Rahman KM, Johura FT, Islam S, Parvin T, Bhuyian MS, Sharif MB, Rahman SR, Sack DA, Sack RB, George CM, Alam M. 2016. Transmission of Infectious Vibrio cholerae through Drinking Water among the Household Contacts of Cholera Patients (CHoBI7 Trial). Front Microbiol 7:1635.

16. Shapiro BJ, Levade I, Kovacikova G, Taylor RK, Almagro-Moreno S. 2016. Origins of pandemic Vibrio cholerae from environmental gene pools. Nat Microbiol 2:16240.

17. Unterweger D, Miyata ST, Bachmann V, Brooks TM, Mullins T, Kostiuk B, Provenzano D, Pukatzki S. 2014. The Vibrio cholerae type VI secretion system employs diverse effector modules for intraspecific competition. Nat Commun 5:3549.

18. Nora Hussein, Paul Kirchberger YFB. Type six secretion mediated competetion in Vibrio cholerae. Unpublished.

19. Islam MT, Liang K, Im MS, Winkjer J, Busby S, Tarr CL, Boucher Y. 2018. Draft Genome Sequences of Nine Vibrio sp. Isolates from across the United States Closely Related to Vibrio cholerae. Microbiol Resour Announc 7.

20. Garrine M, Mandomando I, Vubil D, Nhampossa T, Acacio S, Li S, Paulson JN, Almeida M, Domman D, Thomson NR, Alonso P, Stine OC. 2017. Minimal genetic change in Vibrio cholerae in Mozambique over time: Multilocus variable number tandem repeat analysis and whole genome sequencing. PLoS Negl Trop Dis 11:e0005671.

21. Bishop-Lilly KA, Johnson SL, Verratti K, Luu T, Khiani A, Awosika J, Mokashi VP, Chain PS, Sozhamannan S. 2014. Genome sequencing of 15 clinical Vibrio isolates, including 13 non-o1/non-o139 serogroup strains. Genome Announc 2.

22. Auch AF, von Jan M, Klenk HP, Goker M. 2010. Digital DNA-DNA hybridization for microbial species delineation by means of genome-to-genome sequence comparison. Stand Genomic Sci 2:117–134.

23. Ciufo S, Kannan S, Sharma S, Badretdin A, Clark K, Turner S, Brover S, Schoch CL, Kimchi A, DiCuccio M. 2018. Using average nucleotide identity to improve taxonomic assignments in prokaryotic genomes at the NCBI. Int J Syst Evol Microbiol 68:2386–2392.

24. Richter M, Rossello-Mora R. 2009. Shifting the genomic gold standard for the prokaryotic species definition. Proc Natl Acad Sci U S A 106:19126–19131.

25. Dorman MJ, Kane L, Domman D, Turnbull JD, Cormie C, Fazal MA, Goulding DA, Russell JE, Alexander S, Thomson NR. 2019. The history, genome and biology of NCTC 30: A non-pandemic Vibrio cholerae isolate from World War One. Proc R Soc B Biol Sci https://doi.org/10.1098/rspb.2018.2025.

26. Gardner AD, Venkatraman K V. 1935. The antigens of the cholera group of vibrios. J Hyg (Lond) https://doi.org/10.1017/S0022172400032265.

27. Fiedler G, Pajatsch M, Böck A. 1996. Genetics of a novel starch utilisation pathway present in Klebsiella oxytoca. J Mol Biol https://doi.org/10.1006/jmbi.1996.0085.

28. Davis GH, Park RW. 1962. A taxonomic study of certain bacteria currently classified as Vibrio species. J Gen Microbiol https://doi.org/10.1099/00221287-27-1-101.

29. Sun T, Altenbuchner J. 2010. Characterization of a mannose utilization system in bacillus subtilis. J Bacteriol https://doi.org/10.1128/JB.01673-09.

30. Lee KM, Park Y, Bari W, Yoon MY, Go J, Kim SC, Lee H Il, Yoon SS. 2012. Activation of cholera toxin production by anaerobic respiration of trimethylamine N-oxide in Vibrio cholerae. J Biol Chem https://doi.org/10.1074/jbc.M112.394932.

31. Lin W, Fullner KJ, Clayton R, Sexton JA, Rogers MB, Calia KE, Calderwood SB, Fraser C, Mekalanos JJ. 1999. Identification of a vibrio cholerae RTX toxin gene cluster that is tightly linked to the cholera toxin prophage. Proc Natl Acad Sci U S A https://doi.org/10.1073/pnas.96.3.1071.

32. Dziejman M, Serruto D, Tam VC, Sturtevant D, Diraphat P, Faruque SM, Rahman MH, Heidelberg JF, Decker J, Li L, Montgomery KT, Grills G, Kucherlapati R, Mekalanos JJ. 2005. Genomic characterization of non-O1, non-O139 Vibrio cholerae reveals genes for a type III secretion system. Proc Natl Acad Sci U S A https://doi.org/10.1073/pnas.0409918102.

33. Mutreja A, Kim DW, Thomson N, Connor TR, Hee J, Lebens M, Niyogi SK, Kim EJ, Ramamurthy T, Chun J, Parkhill J, Dougan G. 2013. Evidence for multiple waves of global transmission within the seventh cholera pandemic. Nature https://doi.org/10.1038/nature10392.Evidence.

34. Lugo MR, Merrill AR. 2015. The father, son and cholix toxin: The third member of the DT group mono-ADP-ribosyltransferase toxin family. Toxins (Basel).

35. Choi SY, Rashed SM, Hasan NA, Alam M, Islam T, Sadique A, Johura FT, Eppinger M, Ravel J, Huq A, Cravioto A, Colwell RR. 2016. Phylogenetic diversity of vibrio cholerae associated with endemic cholera in Mexico from 1991 to 2008. MBio https://doi.org/10.1128/mBio.02160-15.

36. Payne SM, Mey AR, Wyckoff EE. 2016. Vibrio Iron Transport: Evolutionary Adaptation to Life in Multiple Environments. Microbiol Mol Biol Rev https://doi.org/10.1128/mmbr.00046-15.

37. Bina XR, Provenzano D, Nguyen N, Bina JE. 2008. Vibrio cholerae RND family efflux systems are required for antimicrobial resistance, optimal virulence factor production, and colonization of the infant mouse small intestine. Infect Immun https://doi.org/10.1128/IAI.01620-07.

38. Trastoy R, Manso T, Fernández-García L, Blasco L, Ambroa A, Pérez Del Molino ML, Bou G, García-Contreras R, Wood TK, Tomás M. 2018. Mechanisms of bacterial tolerance and persistence in the gastrointestinal and respiratory environments. Clin Microbiol Rev.

39. Routh MD, Zalucki Y, Su CC, Long F, Zhang Q, Shafer WM, Yu EW. 2010. Efflux pumps of the Resistance-nodulation-division family: A perspective of their structure, function, and regulation in gram-negative bacteria. Adv Enzymol Relat Areas Mol Biol https://doi.org/10.1002/9780470920541.ch3.

40. Brayton PR, Bode RB, Colwell RR, MacDonell MT, Hall HL, Grimes DJ, West PA, Bryant TN. 1986. Vibrio cincinnatiensis sp. nov., a new human pathogen. J Clin Microbiol https://doi.org/10.1128/jcm.23.1.104-108.1986.

41. Zheng B, Jiang X, Cheng H, Guo L, Zhang J, Xu H, Yu X, Huang C, Ji J, Ying C, Feng Y, Xiao Y, Li L. 2017. Genome characterization of two bile-isolated Vibrio fluvialis strains: An insight into pathogenicity and bile salt adaption. Sci Rep https://doi.org/10.1038/s41598-017-12304-8.

42. Polz MF, Alm EJ, Hanage WP. 2013. Horizontal gene transfer and the evolution of bacterial and archaeal population structure. Trends Genet.

43. Schliep K, Lopez P, Lapointe FJ, Bapteste É. 2011. Harvesting evolutionary signals in a forest of prokaryotic gene trees. Mol Biol Evol https://doi.org/10.1093/molbev/msq323.

44. Wiedenbeck J, Cohan FM. 2011. Origins of bacterial diversity through horizontal genetic transfer and adaptation to new ecological niches. FEMS Microbiol Rev.

45. Faruque SM, Sack DA, Sack RB, Colwell RR, Takeda Y, Nair GB. 2003. Emergence and evolution of Vibrio cholerae O139. Proc Natl Acad Sci U S A https://doi.org/10.1073/pnas.0337468100.

46. Treangen TJ, Ondov BD, Koren S, Phillippy AM. 2014. The harvest suite for rapid core-genome alignment and visualization of thousands of intraspecific microbial genomes. Genome Biol https://doi.org/10.1186/s13059-014-0524-x.

47. Wright JJ, Lee S, Zaikova E, Walsh DA, Hallam SJ. 2009. DNA extraction from 0.22 μM Sterivex filters and cesium chloride density gradient centrifugation. J Vis Exp https://doi.org/10.3791/1352.

48. Alam M, Islam A, Bhuiyan NA, Rahim N, Hossain A, Khan GY, Ahmed D, Watanabe H, Izumiya H, Faruque AS, Akanda AS, Islam S, Sack RB, Huq A, Colwell RR, Cravioto A. 2011. Clonal transmission, dual peak, and off-season cholera in Bangladesh. Infect Ecol Epidemiol 1.

49. Bochner BR, Gadzinski P, Panomitros E. 2001. Phenotype Microarrays for high-throughput phenotypic testing and assay of gene function. Genome Res https://doi.org/10.1101/gr.186501.

50. Kozich JJ, Westcott SL, Baxter NT, Highlander SK, Schloss PD. 2013. Development of a dual-index sequencing strategy and curation pipeline for analyzing amplicon sequence data on the miseq illumina sequencing platform. Appl Environ Microbiol https://doi.org/10.1128/AEM.01043-13.

51. R Development Core Team R. 2011. R: A Language and Environment for Statistical ComputingR Foundation for Statistical Computing.

52. Callahan BJ, McMurdie PJ, Rosen MJ, Han AW, Johnson AJA, Holmes SP. 2016. DADA2: High-resolution sample inference from Illumina amplicon data. Nat Methods https://doi.org/10.1038/nmeth.3869.

53. Schloss PD, Westcott SL, Ryabin T, Hall JR, Hartmann M, Hollister EB, Lesniewski RA, Oakley BB, Parks DH, Robinson CJ, Sahl JW, Stres B, Thallinger GG, Van Horn DJ, Weber CF. 2009. Introducing mothur: Open-source, platform-independent, community-supported software for describing and comparing microbial communities. Appl Environ Microbiol https://doi.org/10.1128/AEM.01541-09.

54. Oksanen J, Kindt R, Legendre P, O’Hara B, Simpson GL, Solymos PM, Stevens MHH, & Wagner H. 2008. The vegan package. Community Ecol Packag.

55. Angiuoli S V., Salzberg SL. 2011. Mugsy: Fast multiple alignment of closely related whole genomes. Bioinformatics https://doi.org/10.1093/bioinformatics/btq665.

56. Stamatakis A. 2014. RAxML version 8: A tool for phylogenetic analysis and post-analysis of large phylogenies. Bioinformatics https://doi.org/10.1093/bioinformatics/btu033.

57. Aziz RK, Bartels D, Best A, DeJongh M, Disz T, Edwards RA, Formsma K, Gerdes S, Glass EM, Kubal M, Meyer F, Olsen GJ, Olson R, Osterman AL, Overbeek RA, McNeil LK, Paarmann D, Paczian T, Parrello B, Pusch GD, Reich C, Stevens R, Vassieva O, Vonstein V, Wilke A, Zagnitko O. 2008. The RAST Server: Rapid annotations using subsystems technology. BMC Genomics https://doi.org/10.1186/1471-2164-9-75.

58. Kearse M, Moir R, Wilson A, Stones-Havas S, Cheung M, Sturrock S, Buxton S, Cooper A, Markowitz S, Duran C, Thierer T, Ashton B, Meintjes P, Drummond A. 2012. Geneious Basic: An integrated and extendable desktop software platform for the organization and analysis of sequence data. Bioinformatics https://doi.org/10.1093/bioinformatics/bts199.

59. Chaudhari NM, Gupta VK, Dutta C. 2016. BPGA-an ultra-fast pan-genome analysis pipeline. Sci Rep https://doi.org/10.1038/srep24373.

