## Supplemental File for "Population analysis of *Vibrio cholerae* in aquatic reservoirs reveals a novel sister species *(Vibrio paracholerae* sp. nov.) with a history of association with human infections"

**Supplementary material**

**Description of the novel species *Vibrio paracholerae***

*Vibrio paracholerae* (pa.ra.chol.er.ae. Gr. prep. para alongside; L. gen. f. n. cholerae bilious disease; referring to the isolation of the type strain alongside *Vibrio cholerae*, the causative agent of cholera.

A Gram stain negative, oxidase positive, curved rod-shaped bacterium, roughly 1.25-2 mm in length and 0.4 mm in width. Exhibits motility by means of a single polar flagellum. The ability to utilize α-cyclodextrin and pectin as well as the lack of ability to utilize mannose, L-aspartic acid and citric acid differentiates the species from its closest relative *V. cholerae* (99% identity of 16S *rRNA* gene). Forms *V. cholerae*-like yellow circular colonies on TCBS agar and circular colonies of creamy-white colour on TSB agar. Positive for carbon utilization from D-glucose, D-fructose, sucrose, maltose, D-galactose, maltotriose, D-mannitol, D-trehalose, D glucose-6-phosphate, N-acetyl-D-glucosamine, glycerol, succinic acid, L-Lactic Acid, L-glutamic acid, fumar acid, acetic acid, L-proline, D-alanine, L-asparagine, 2-deoxy adenosine, adenosine, inosine and L-serine, dextrin, gelatin, glycogen, D-glucosamine and D-lactic acid methyl ester.  Variation of response observed between strains in utilization of D-mannose (2/6), D-cellobiose, D-fructose-6-phosphate, D-psicose, α-D-lactose, L-aspartic acid, D-glucuronic Acid, D-gluconic acid, D,L-α-glycerol phosphate, D,L-malic acid, D-ribose, tween 20, thymidine, α-keto-glutaric acid, tween 40, tween 80, α-keto-butyric acid, uridine, L-glutamine, α-hydroxy-glutaric acid-γ-lactone, ß-methyl-D-glucoside, citric acid, L-threonine, L-alanine, L-alanyl-glycine, glycine , histidine, methyl pyruvate, D-malic acid, glycyl-L-proline and pyruvic acid.

The proposed type strain EDC-792 was isolated from environmental water in Dhaka, Bangladesh in 2016; the strain displays all the properties given above for the species.

**Supplementary tables and figures:**

| Location | Number of samples | Salinity range (ppt) | DO range (mg/L) | pH range | Temperature range (°C) |
|---|---|---|---|---|---|
| Dhaka | 60 | 0-0.8 | 0.19-5.05 | 6.4-7.7 | 27.1-33.6 |
| Oyster Pond | 40 | 0-6 | 5.4-8.6 | 5.5-7.5 | 10.5-28 |

**Table S1: Physiochemical data ranges of the two sampling locations used in the study.**

| **Isolates** | **Origin** | **Year of isolation** | **Source** | **Phylogenetic group** | **Genome accession number** |
|---|---|---|---|---|---|
| N16961 | Bangladesh | 1971 | Clin | PG *Vibrio cholerae* O1 | GCA_000006745.1 |
| 2010EL 1786 | Haiti | 2010 | Clin | PG *Vibrio cholerae* O1 | GCA_000166455.2 |
| EDC 721 | Bangladesh | 2015 | Env | PG *Vibrio cholerae* O1 | WYCI00000000 |
| MJ 1236 | Bangladesh | 1994 | Clin | PG *Vibrio cholerae* O1 | GCA_000022585.1 |
| MO10 | India | 1992 | Clin | PG *Vibrio cholerae* O1 | GCA_000152425.1 |
| 2740-80 | USA | 1980 | Env | PG *Vibrio cholerae* O1 | GCA_001683415.1 |
| BX3330286 | Australia | 1985 | Env | PG *Vibrio cholerae* O1 | GCA_000174335.1 |
| O395 | India | 1965 | Clin | PG *Vibrio cholerae* O1 | GCA_000021625.1 |
| 95412 | Mexico | 1997 | Env | PG *Vibrio cholerae* O1 | GCA_000348105.2 |
| V52 | Sudan | 1968 | Clin | PG *Vibrio cholerae* O37 | GCA_000167935.2 |
| 2012EL1759 | Haiti | 2012 | Env | PG Vibrio cholerae | JNEW01000000 |

| | | | | | |
|---|---|---|---|---|---|
| | | | | O1 | |
| 2012Env9 | Haiti | 2012 | Env | Non-PG *Vibrio cholerae* O1 | GCA_000788715.2 |
| MZO-03 | Bangladesh | 2001 | Clin | Non-PG *Vibrio cholerae* non-O1/O139 | GCA_000168935.3 |
| 12129 | Australia | 1985 | Env | Non-PG *Vibrio cholerae* O1 | ACFQ00000000 |
| 1587 | Peru | 2000 | Clin | Non-PG *Vibrio cholera*e non-O1/O139 | GCA_000168895.2 |
| CP1035 | Mexico | 2004 | Clin | Non-PG *Vibrio cholerae* non-O1/O139 | AJRM00000000 |
| EDC 689 | Bangladesh | 2015 | Env | Non-PG *Vibrio cholerae* non-O1/O139 | WYCR00000000 |

| | | | | | |
|---|---|---|---|---|---|
| EDC 715 | Bangladesh | 2016 | Env | Non-PG *Vibrio cholerae* non-O1/O139 | WYCJ00000000 |
| EDC 800 | Bangladesh | 2016 | Env | Non-PG *Vibrio cholerae* non-O1/O139 | WYCA00000000 |
| EM1676A | Bangladesh | 2011 | Env | Non-PG *Vibrio cholerae* non-O1/O139 | GCA_000348345.2 |
| 2012ENV-92 | Haiti | 2012 | Env | Non-PG *Vibrio cholerae* non-O1/O139 | GCA_000788755.1 |
| HE 48 | Haiti | 2010 | Env | Non-PG *Vibrio cholerae* non-O1/O139 | GCA_000220785.2 |
| VCC19 | Brazil | 1994 | Env | *Vibrio paracholerae* | GCA_000438805.2 |
| 877163 | Bangladesh | 2002 | Env | *Vibrio paracholerae* | GCA_001402745.1 |
| EDC 792 | Bangladesh | 2016 | Env | *Vibrio paracholerae* | WYCC00000000 |

| | | | | | |
|---|---|---|---|---|---|
| EDC 690 | Bangladesh | 2015 | Env | *Vibrio paracholerae* | WUWI00000000 |
| EDC 716 | Bangladesh | 2015 | Env | *Vibrio paracholerae* | WYBZ00000000 |
| EDC 717 | Bangladesh | 2015 | Env | *Vibrio paracholerae* | WYBY00000000 |
| HE09 | Haiti | 2010 | Env | *Vibrio paracholerae* | GCA_000221405.1 |
| HE16 | Haiti | 2010 | Env | *Vibrio paracholerae* | GCA_000303085.1 |
| CISM300506 | Mozambique | 2008 | Clin | *Vibrio paracholerae* | GCA_002097755.1 |
| CISM1163068 | Mozambique | 2012 | Clin | *Vibrio paracholerae* | GCA_002097815.1 |
| 49093 DA89 | Thailand | 1992 | Clin | *Vibrio paracholerae* | GCA_000737015.1 |
| SIO | USA | 2003 | Waste water | *Vibrio paracholerae* | GCA_001857455.1 |
| 2017V 1144 | USA | 2017 | Stool | *Vibrio paracholerae* | GCA_003312015.1 |
| 2014V 1107 | USA | 2014 | Stool | *Vibrio paracholerae* | GCA_003311945.1 |
| 2017V 1105 | USA | 2017 | Wound | *Vibrio paracholerae* | GCA_003311975.1 |
| 2016V 1114 | USA | 2016 | Stool | *Vibrio paracholerae* | GCA_003312085.1 |
| 2016V 1111 | USA | 2016 | Stool | *Vibrio paracholerae* | GCA_003311965.1 |
| 2017V 1176 | USA | 2017 | Animal feed | *Vibrio paracholerae* | GCA_003312095.1 |
| 2016V 1091 | USA | 2016 | Stool | *Vibrio paracholerae* | GCA_003312065.1 |
| 2017V 1110 | USA | 2017 | Wound | *Vibrio paracholerae* | GCA_003312005.1 |
| 87395 | UK | UK | UK | *Vibrio paracholerae* | GCA_000348085.2 |
| 07 2425 | UK | UK | UK | *Vibrio paracholerae* | GCA_003311905.1 |
| NCTC30 | Egypt | 1916 | Clin | *Vibrio paracholerae* | LS997868.1 |

**Table S2: Demographic information of the *V. paracholerae* sp. nov. and *V. cholerae* isolates used in this study.** Clin: clinical; Env: environmental; UK: Unknown.

| **Chemical/ heavy metal** | ***Vibrio paracholerae* sp. nov.** | | | | ***Vibrio cholerae*** | | | |
|---|---|---|---|---|---|---|---|---|
| | **EDC 690** | **EDC 716** | **EDC 792** | **2016V-1091** | **N16961** | **V52** | **YB3B05** | **YB8E08** |
| Cadmium chloride | R | R | R | ND | S | S | ND | ND |
| Sodium selenite | R | R | R | ND | S | S | ND | ND |
| Dichlofuanid | R | R | R | ND | S | S | ND | ND |

**Table S3: Responses towards chemical/heavy metals of *Vibrio paracholerae* sp. nov., differentiating it from its closest relatives *Vibrio cholerae*.** R: resistant; S: sensitive; ND: not done.

| Strain | Phylogenetic group | CTX-VPI1-VPI2 | RTX toxin | SXT element | T3SS | RND efflux gene cluster | Cholix toxin (chxA) | β-lactamase |
|---|---|---|---|---|---|---|---|---|
| N16961 | *V. cholerae* O1 El Tor (PG) | + | + | - | - | - | - | - |
| O395 | *V. cholerae* O1 classical (PG) | + | + | - | - | - | - | - |
| MO10 | *V. cholerae* O139 (PG) | + | + | + | - | - | - | - |
| 12129 | *V. cholerae* non-O1/O139 (non-PG) | - | + | - | + | - | - | - |
| VCC19 | *V. paracholerae* sp. nov. | - | + | + | - | + | - | + |
| 877163 | | - | + | - | + | + | + | - |
| EDC-792 | | - | + | - | - | + | + | - |
| EDC-690 | | - | + | - | - | - | + | - |
| EDC-716 | | - | + | - | - | + | - | - |
| EDC-717 | | - | + | - | - | + | - | - |
| HE09 | | - | + | - | - | + | + | - |
| HE16 | | - | + | - | - | + | - | - |
| CISM300506 | | - | + | + | - | + | + | + |
| CISM1163068 | | - | + | + | - | + | + | + |
| 49093-DA89 | | - | + | - | - | + | - | + |
| SIO | | - | + | - | - | - | + | - |
| 2017V-1144 | | - | + | - | - | + | - | + |
| 2014V-1107 | | - | + | + | - | + | + | - |
| 2017V-1105 | | - | + | + | - | + | + | + |

| | | | | | | | | |
|---|---|---|---|---|---|---|---|---|
| 2016V-1114 | | - | + | - | + | + | - | - |
| 2016V-1111 | | - | + | - | + | + | - | - |
| 2017V-1176 | | - | + | - | - | + | - | + |
| 2016V-1091 | | - | + | - | - | + | + | - |
| 2017V-1110 | | - | + | - | - | - | - | - |
| 87395 | | - | + | - | - | + | + | + |
| 07-2425 | | - | - | - | - | + | - | - |
| NCTC 30 | | - | + | - | + | + | - | + |

**Table S4: Presence of potential virulence traits in *V. paracholerae* sp. nov isolates in comparison to *V. cholerae* reference strains**

| Strain | Origin | Source | dDDH with *V. paracholerae* sp. nov (EDC792*)* | NCBI genome accession number |
|---|---|---|---|---|
| N2784 | China | Clin | 85.2 | VSHN01000030.1 |
| N2770 | China | Clin | 85.7 | VSHB01000078.1 |
| N2768 | China | Clin | 89.4 | VSGZ01000037.1 |
| EL2338 | China | Clin | 89.1 | VMPB01000147.1 |
| 2204 | Brazil | Env | 85.8 | VHOF01000002.1 |
| 2290 | Brazil | Env | 79.1 | VHOE01000001.1 |
| N2748 | China | Clin | 83.7 | VSGI01000001.1 |
| EL2403 | China | Clin | 83.2 | VMOL01000089.1 |
| A110912Z3 | Austria | Env | 82.4 | VIQC01000020.1 |
| N2807 | China | Clin | 83 | VSID01000044.1 |
| N2795 | China | Clin | 83.2 | VSHY01000044.1 |
| N2794 | China | Clin | 83.4 | VSHX01000029.1 |
| N2791 | China | Clin | 83.3 | VSHU01000058.1 |
| A3_296 | Brazil | Clin | 80.6 | QBJE01000015.1 |
| FORC_076 | South korea | Clin | 86 | NZ_CP026531 |

**Table S5: Information of genomes falling into the *V. paracholerae* sp. nov clade disclosed in the NCBI database after 2019.** dDDH: digital DNA-DNA hybridization value; Clin: clinical; Env: environmental.
